# Cxcr2 is required for ductal outgrowth and shapes the stromal microenvironment in postnatal mammary gland development

**DOI:** 10.64898/2026.11.08.765094

**Authors:** Courtney L. Flatt, Emma M. Powers, Aaron Rosenberg, Meghan O’Hare, Anh Bui, Ann Dharshika Shangaradas, Junmin Wu, Glen L. Niebur, Jun Li, Laurie E. Littlepage

## Abstract

Mammary gland development requires coordinated signaling between epithelial and stromal compartments. While CXC chemokine receptor 2 (CXCR2) is known to regulate inflammatory and tissue remodeling processes in disease, including breast cancer, its role in mammary gland morphogenesis is not well understood. Here, we examined postnatal mammary gland development in germline Cxcr2 knockout mice. Loss of Cxcr2 resulted in delayed ductal outgrowth, reduced epithelial area, and altered stromal architecture, including decreased adipocyte size and increased collagen deposition, despite preserved branching organization. These defects occurred in the context of severe systemic immune dysregulation, marked by expansion of granulocytic populations and loss of adaptive immune cells. Together, our findings indicate that CXCR2 indirectly regulates mammary gland development through coordination between the epithelium and stroma, rather than acting through epithelial cell-intrinsic mechanisms. Because the same myeloid expansion, immune suppression, and collagen-rich stromal remodeling that CXCR2 controls during normal development are hallmarks of the pro-tumorigenic breast microenvironment, these findings establish CXCR2 as a regulator of the immune and stromal landscape of the mammary gland and provide a developmental framework for understanding its contributions to the breast tumor microenvironment.

## Introduction

Development of the mammary gland requires a highly regulated interplay between epithelial cells and the surrounding stroma, including adipocytes, endothelial cells, fibroblasts, and immune cells [1]. During postnatal development, coordinated signaling between these cell types drives ductal elongation, epithelial branching morphogenesis, angiogenesis, and extracellular matrix remodeling of the mammary gland. Chemokines, or chemotactic cytokines involved in immune cell recruitment, are increasingly recognized as key mediators of epithelial-stromal crosstalk, coordinating cell trafficking, stromal remodeling, and paracrine signaling between immune, adipose, and epithelial cells during mammary development [2–5].

CXC chemokine receptor 2 (CXCR2; mouse: Cxcr2) is a G-protein-coupled receptor for ELR+ CXC chemokines [6] and is an essential regulator of neutrophil maturation and mobilization from bone marrow [7]. Beyond its role in innate immunity, CXCR2 promotes angiogenesis and regulates cell migration and inflammatory responses during infection and wound healing [8–10]. Emerging evidence further implicates CXCR2 in cancer progression, including breast cancer, where CXCR2 expression correlates with increased tumor growth and metastasis and with poor prognosis [11–16]. Notably, CXCL5/CXCR2 signaling promotes breast cancer colonization in bone, and osteoblasts have been identified as a source of CXCL5 that supports tumor progression in bone mimetic models [16, 17]. In these settings, CXCR2 coordinates the immune and stromal microenvironment to support tumor progression. Whether CXCR2 governs these compartments during normal mammary development, and how that role relates to its established functions in the tumor microenvironment, remains incompletely defined.

Cxcr2 functions in multiple stromal cells relevant to mammary gland development, including adipocytes and immune cells. Cxcr2 signaling induces mesenchymal stem cell differentiation to adipocytes through macrophage-derived Cxcl1 [18]. Adipocyte differentiation decreases in cell culture with Cxcr2 antagonist treatment, and adipogenesis-associated genes decrease in the skin and inguinal adipose tissues after Cxcr2 deficiency [19]. In addition, systemic alterations in Cxcr2 KO (BALB/c) mice, including defects in pituitary hormone signaling, were linked to reproductive abnormalities in the mammary gland and ovary [20]. Importantly, these defects were rescued when mice were housed in specific and opportunistic pathogen-free conditions, underscoring the contribution of chronic inflammation to the Cxcr2 KO phenotype. Together, these findings suggest that CXCR2 may influence mammary gland development through systemic, stromal, or immune-mediated mechanisms.

Puberty represents a critical window of mammary gland development, during which the rudimentary epithelial ductal tree undergoes rapid expansion and branching to establish the gland’s functional architecture. This process depends on finely tuned interactions between epithelial cells and the surrounding stroma, including immune cells and extracellular matrix. Despite established roles for CXCR2 signaling in tissue remodeling and in epithelial-stromal interactions in cancer, its involvement in orchestrating and regulating normal mammary gland development, particularly during puberty, has not been studied.

In this study, we investigated the role of Cxcr2 in postnatal mammary gland development and immune homeostasis using Cxcr2 wild-type (WT), heterozygous (Het), and knockout (KO) mice on the FVB/N background. By integrating immune profiling, mammary gland whole-mount analysis, histology, and transplantation approaches, we sought to determine whether Cxcr2 regulates mammary branching morphogenesis through epithelial cell-intrinsic or epithelial-stromal signaling.

## Results

### Cxcr2 KO mice on FVB/N background have decreased survival compared to WT mice

We previously reported that germline Cxcr2 deletion on the FVB/N background results in developmental defects and decreased survival [21]. Here, using a substantially expanded cohort (N=3163), we confirm that Cxcr2 KO FVB mice show significantly decreased survival compared to Cxcr2 WT and Het mice (P<0.0001; Figure 1A), and re-examine these systemic phenotypes as context for the mammary gland analysis that follows.

**Fig. 1.**
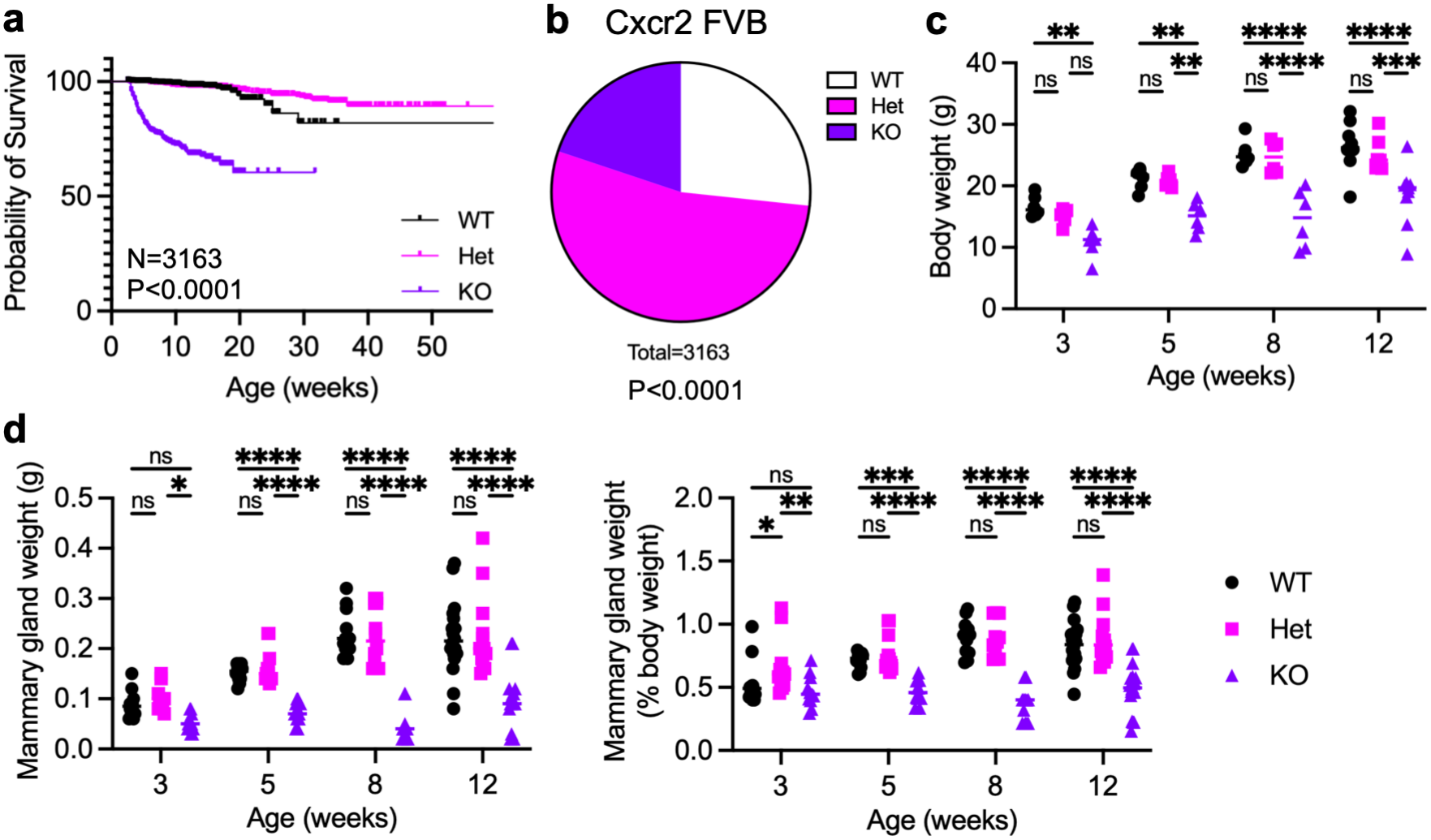
Cxcr2 KO mice show decreased survival, reduced body weight, and reduced mammary gland weight. (A) Kaplan-Meier survival analysis of Cxcr2 WT, Het, and KO mice; significance by logrank (Mantel-Cox) test (n = 844 WT, 1690 Het, and 629 KO; 3163 total mice). (B) Genotype distribution of Cxcr2 WT, Het, and KO mice from Het x Het crosses compared to expected Mendelian ratios by chi-square test (n = 3163 mice). (C) Body weight (g) of Cxcr2 WT, Het, and KO virgin female mice at 3, 5, 8, and 12 weeks of age (n = 6-9/genotype/age). (D) Mammary gland weight in grams (left) and as a percentage of total body weight (right) at 3, 5, 8, and 12 weeks. Statistical analysis by two-way ANOVA with Tukey’s multiple comparisons test. Lines indicate mean. ns = not significant (p ≥ 0.05), *p < 0.05, **p < 0.01, ***p < 0.001, ****p < 0.0001. Expanded cohort including data previously reported in [21]

We assessed the frequency of Cxcr2 WT, Het, and KO genotypes in Cxcr2 Het x Het offspring and compared them to expected ratios for Mendelian inheritance (Figure 1B, Supplemental Figure 1A). Across 3163 total mice, the genotype ratios differed significantly from expected (P<0.0001); 53.4% were Het mice, while only 19.9% were KO, and 26.7% were WT mice. In female mice, only 19.7% were Cxcr2 KO, while 26.5% were WT (P<0.0001). This shift was not statistically significant in male mice (27.4% WT, 51.6% Het, 21.0% KO; n=537, P=0.0888).

### Cxcr2 KO mice are smaller and have less body fat than littermates

We measured body weight in 3-, 5-, 8-, and 12-week-old virgin female mice (Figure 1C; n=6/genotype). Body composition at 3, 5, and 8 weeks was previously reported [21]; here we extend this analysis to 12 weeks to capture the full pubertal-to-postpubertal window. The Cxcr2 KO mice were significantly smaller than WT and Het littermates from birth and had delayed growth up to 12 weeks. The mean body weight in 3-week-old Cxcr2 KO mice was only 10.85 g, compared to 16.67 g in 3-week-old WT mice. At 5 weeks, Cxcr2 KO mean body weight was 15.07 g, versus 21.18 g for WT, and at 8 weeks, the average weight was 14.64 g versus 25.28 g in Cxcr2 KO and WT mice, respectively. By 12 weeks of age, the mean body weight was 26.31 g in WT and 18.53 g in Cxcr2 KO mice. Thus, by 8 weeks of age, female Cxcr2 KO mice were only 58% of the normal weight of their WT littermates (P<0.0001) and by 12 weeks were 70% that of WT mice (P=0.0008). Therefore, Cxcr2 deficiency significantly limited the ability of these mice to grow and thrive.

To assess body fat and lean content, we used EchoMRI body composition analysis in 3-, 5-, 8-, and 12-week-old female mice (Supplemental Figure 1B; n=6/genotype). As previously reported [21] and extended here to 12 weeks, body fat content (as a percentage of body weight) in female Cxcr2 KO mice was markedly decreased after three weeks old, with a 40% reduction in 5-week-old mice, a 55% decrease in 8-week-old mice, and 40% reduction in 12-week-old mice. Accordingly, lean content did not differ significantly in 3- or 5-week-old mice but increased 14% and 12%, respectively, in 8-week-old and 12-week-old KO mice compared to WT littermates.

To examine the effects of Cxcr2 deficiency on the mammary gland, we collected the fourth abdominal mammary glands from estrus-matched 3-, 5-, 8-, and 12-week-old virgin mice. Mammary gland weights were significantly less in Cxcr2 KO mice, both in mass and as a percent of total body weight (Figure 1D). The mean mammary gland weight in 3-week-old Cxcr2 KO mice was only 0.053 g, compared to 0.088 g in 3-week-old WT mice. At 5 weeks, the Cxcr2 KO mean mammary gland weight was 0.073 g versus 0.152 g for WT, while at 8 weeks old the mean mammary gland weight was 0.046 g versus 0.230 g in Cxcr2 KO and WT mice, respectively. Thus, by 8 weeks of age, Cxcr2 KO mammary glands were only 20% of the normal weight compared to their WT littermates (P<0.0001).

We accounted for the reduced body weight of the KO animals by determining mammary gland weight as a percentage of body weight. By 8 weeks of age, WT mean mammary glands weights were 0.89%, versus only 0.36% in Cxcr2 KO mice, which is a 60% reduction in normalized mammary gland weight (P<0.0001).

### Splenomegaly and enlarged cervical lymph nodes in Cxcr2 KO mice

We observed gross abnormalities in spleens and cervical lymph node sizes when compared between genotypes (Figure 2A-B). Spleens of Cxcr2 KO mice were enlarged to three times the normal size, with a mean weight of 0.347 g in KO compared to 0.114 g in WT (Figure 2A). Cervical lymph nodes were enlarged in weight and relative size compared to body weight in Cxcr2 KO mice (Figure 2B). These alterations suggested potential immunological alterations in the Cxcr2 KO mice. This could be due to vivarium pathogens. Our pathogen screening identified the presence of several pathogens, including *Helicobacter ganmani*, *Rodentibacter heylii*, and *Entamoeba muris*, in both WT and Cxcr2 KO animals, suggesting that pathogen-induced inflammation could contribute to the observed proinflammatory state [21].

**Fig. 2.**
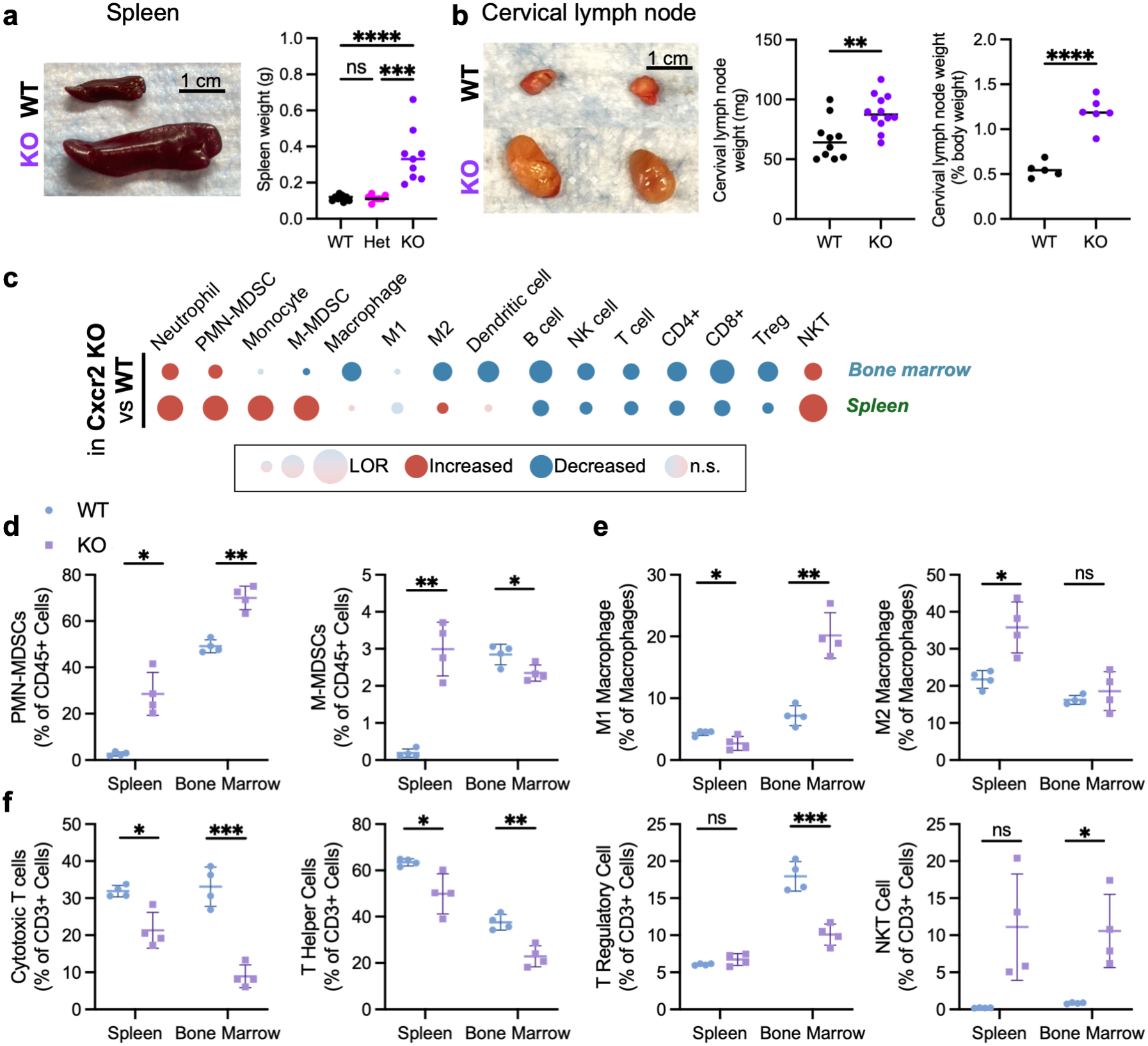
**Cxcr2 deficiency reshapes the splenic and bone marrow immune landscape toward an immunosuppressive, innate-skewed profile. (**A) Representative images and quantification of spleen weight (g) in Cxcr2 WT, Het, and KO mice; scale bar = 1 cm. (B) Representative images and quantification of cervical lymph node weight (mg, left) and as a percentage of body weight (right) in Cxcr2 WT and KO mice; scale bar = 1 cm. (C) Summary of immune cell population changes in Cxcr2 KO versus WT spleen and bone marrow by flow cytometry. Dot color indicates direction of change (red, increased; blue, decreased; pale, not significant) and dot size indicates log odds ratio (LOR). (D–F) Flow cytometry quantification in spleen and bone marrow of Cxcr2 WT and KO mice (n = 4/genotype). (D) PMN-MDSCs and M-MDSCs (as % of CD45+ cells), (E) M1 and M2 macrophages (as % of total macrophages), and (F) cytotoxic T cells, T helper cells, regulatory T cells, and NKT cells (as % of CD3+ cells). Statistical analysis of spleen weight by ANOVA, lymph node weight and immune cell populations by Welch’s t-test. Lines indicate mean. ns, not significant; *p < 0.05, **p < 0.01, ***p < 0.001, ****p < 0.0001

### Cxcr2 deficiency skews the splenic and bone marrow immune compartment toward suppressive myeloid populations

We previously characterized the systemic immune phenotype of Cxcr2 KO FVB/N mice by complete blood count with differential and serum cytokine profiling, which revealed a marked expansion of circulating neutrophils, a reduced lymphocyte fraction, and broad elevation of pro-inflammatory cytokines including G-CSF, CXCL1, CXCL5, and IL-17 [21]. These analyses established that Cxcr2 loss drives granulocyte expansion but did not resolve the specific immune populations underlying this phenotype.

To define the immune landscape at higher resolution and to determine whether Cxcr2 deficiency reshapes the lymphoid organ immune compartment, we profiled the CD45+ immune cell populations in spleen and bone marrow of Cxcr2 WT and KO mice by multiparameter flow cytometry (Figure 2C, Supplemental Figure 2, Table 1-2). Compared to in spleens from WT mice, several myeloid cell populations were altered by Cxcr2 deficiency. Granulocytic and suppressive myeloid populations, including Ly6G+ neutrophils and both PMN-MDSCs (CD11b+ Ly6G+ Ly6C^int^ Arginase1+(Arg1+)) and M-MDSCs (CD11b+ Ly6G-Ly6C^hi^ Arg1+), increased in Cxcr2 KO mice compared to in WT mice (Figure 2D). Monocytic cells (T/B/NK-Ly6G-) and classical monocytes (Ly6C^hi^ Ly6G-) increased with Cxcr2 deficiency. While total macrophages (F480+ Ly6C^hi^ Ly6G-) as percentage of CD45+ cells were unchanged based on Cxcr2 status, their subtype distribution changed. Within the macrophage compartment, Arg1+ M2 macrophages increased, while CD86+ M1 macrophages decreased, consistent with a shift toward a more M2-skewed macrophage profile in the spleen with Cxcr2 deficiency (Figure 2E).

**Table 1.**
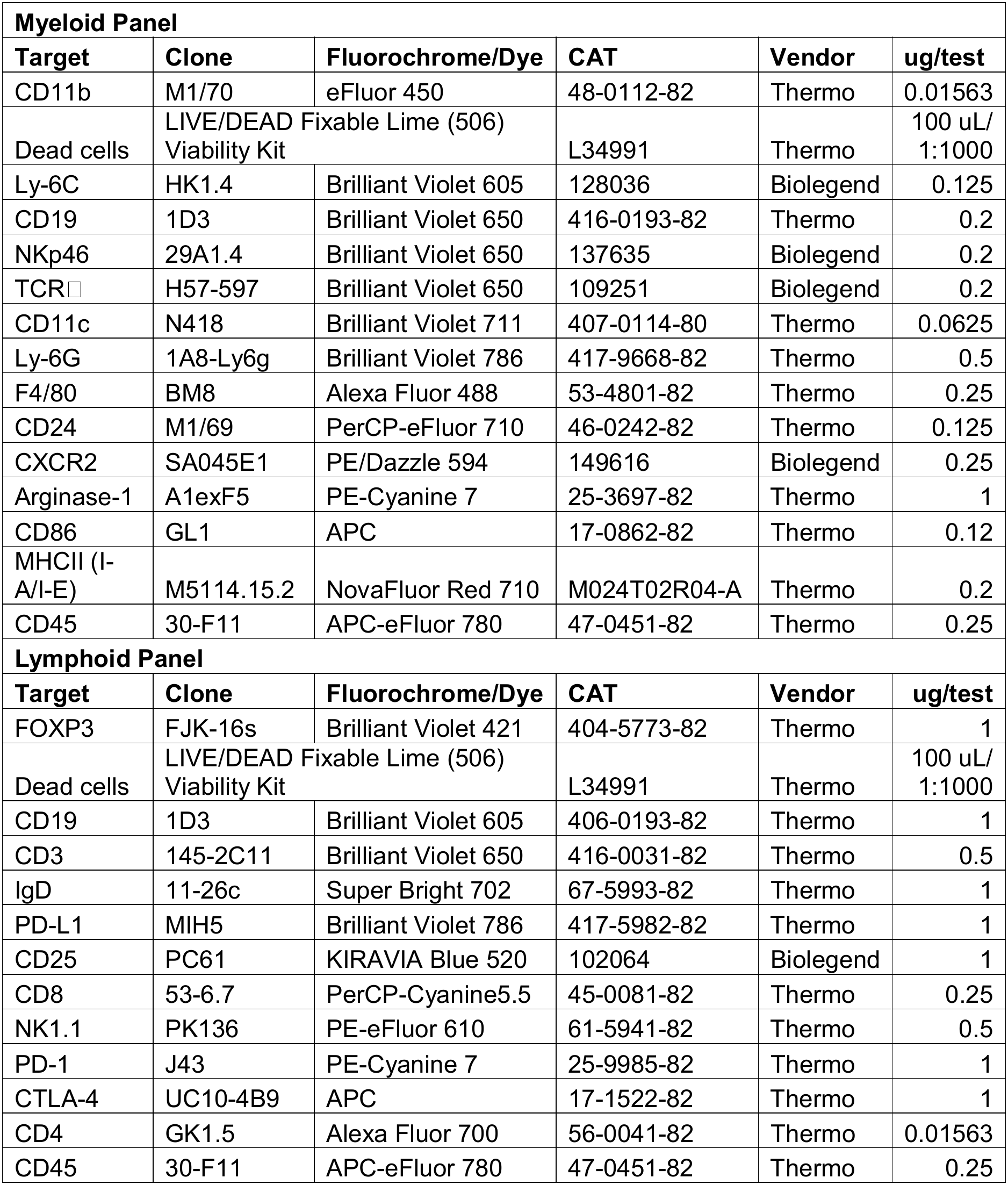
Primary antibodies used for flow cytometry staining. Antibody targets, conjugates, catalog number, and working concentration for spleen and bone marrow immunophenotyping.

| **Myeloid Panel** |  |  |  |  |  |
|---|---|---|---|---|---|
| **Target** | **Clone** | **Fluorochrome/Dye** | **CAT** | **Vendor** | **ug/test** |
| CD11b | M1/70 | eFluor 450 | 48-0112-82 | Thermo | 0.01563 |
| Dead cells | LIVE/DEAD Fixable Lime (506) Viability Kit |  | L34991 | Thermo | 100 uL/ 1:1000 |
| Ly-6C | HK1.4 | Brilliant Violet 605 | 128036 | Biolegend | 0.125 |
| CD19 | 1D3 | Brilliant Violet 650 | 416-0193-82 | Thermo | 0.2 |
| NKp46 | 29A1.4 | Brilliant Violet 650 | 137635 | Biolegend | 0.2 |
| TCR□ | H57-597 | Brilliant Violet 650 | 109251 | Biolegend | 0.2 |
| CD11c | N418 | Brilliant Violet 711 | 407-0114-80 | Thermo | 0.0625 |
| Ly-6G | 1A8-Ly6g | Brilliant Violet 786 | 417-9668-82 | Thermo | 0.5 |
| F4/80 | BM8 | Alexa Fluor 488 | 53-4801-82 | Thermo | 0.25 |
| CD24 | M1/69 | PerCP-eFluor 710 | 46-0242-82 | Thermo | 0.125 |
| CXCR2 | SA045E1 | PE/Dazzle 594 | 149616 | Biolegend | 0.25 |
| Arginase-1 | A1exF5 | PE-Cyanine 7 | 25-3697-82 | Thermo | 1 |
| CD86 | GL1 | APC | 17-0862-82 | Thermo | 0.12 |
| MHCII (I-A/I-E) | M5114.15.2 | NovaFluor Red 710 | M024T02R04-A | Thermo | 0.2 |
| CD45 | 30-F11 | APC-eFluor 780 | 47-0451-82 | Thermo | 0.25 |
| **Lymphoid Panel** |  |  |  |  |  |
| **Target** | **Clone** | **Fluorochrome/Dye** | **CAT** | **Vendor** | **ug/test** |
| FOXP3 | FJK-16s | Brilliant Violet 421 | 404-5773-82 | Thermo | 1 |
| Dead cells | LIVE/DEAD Fixable Lime (506) Viability Kit |  | L34991 | Thermo | 100 uL/ 1:1000 |
| CD19 | 1D3 | Brilliant Violet 605 | 406-0193-82 | Thermo | 1 |
| CD3 | 145-2C11 | Brilliant Violet 650 | 416-0031-82 | Thermo | 0.5 |
| IgD | 11-26c | Super Bright 702 | 67-5993-82 | Thermo | 1 |
| PD-L1 | MIH5 | Brilliant Violet 786 | 417-5982-82 | Thermo | 1 |
| CD25 | PC61 | KIRAVIA Blue 520 | 102064 | Biolegend | 1 |
| CD8 | 53-6.7 | PerCP-Cyanine5.5 | 45-0081-82 | Thermo | 0.25 |
| NK1.1 | PK136 | PE-eFluor 610 | 61-5941-82 | Thermo | 0.5 |
| PD-1 | J43 | PE-Cyanine 7 | 25-9985-82 | Thermo | 1 |
| CTLA-4 | UC10-4B9 | APC | 17-1522-82 | Thermo | 1 |
| CD4 | GK1.5 | Alexa Fluor 700 | 56-0041-82 | Thermo | 0.01563 |
| CD45 | 30-F11 | APC-eFluor 780 | 47-0451-82 | Thermo | 0.25 |

**Table 2.** Cell markers for immunophenotyping. Definition of cell types analyzed by flow cytometry for lymphoid and myeloid panels.

| **Myeloid** |  |
|---|---|
| Leukocyte | CD45+ |
| Myeloid Cell | CD45+ T,B,NK- |
| T,B, or NK cell | CD45+ T,B,NK+ |
| M MDSC | CD45+ CD11b+ Arginase1+ Ly-6Chi Ly-6G- |
| PMN MDSC | CD45+ CD11b+ Arginase1+ Ly-6Cint Ly-6G+ |
| Neutrophil | CD45+ Ly-6G+ |
| Classical Monocytes | CD45+ Ly-6G- Ly-6Chi CD11b+ |
| Dendritic Cells | CD45+ Ly-6G- non-(Ly6Chi CD11b+) CD24+ CD11c+ MHCII+ |
| Macrophage | CD45+ Ly-6G- non-(Ly6Chi CD11b+) F4 80+ |
| M1 Macrophage CD86+ F4 80+ | CD45+ Ly-6G- non-(Ly6Chi CD11b+) F4 80+ CD86+ |
| M2 Macrophage Arg1+ F4 80+ | CD45+ Ly-6G- non-(Ly6Chi CD11b+) F4 80+ Arginase1+ |
| **Lymphoid** |  |
| Leukocyte | CD45+ |
| B Cells | CD45+ CD19+ |
| Naive Unswitched B cell | CD45+ CD19+ IgD+ |
| T Cell | CD45+ CD3+ |
| Cytotoxic T Cell | CD45+ CD3+ CD8+ |
| T Helper Cell | CD45+ CD3+ CD4+ |
| Treg | CD45+ CD3+ CD4+ FoxP3+ CD25+ |
| NK Cells | CD45+ NK1.1+ CD3- |
| NKT Cell | CD45+ NK1.1+ CD3+ |

Lymphoid lineage cell populations also were affected by Cxcr2. The decreased CD45+ cell populations in the Cxcr2 KO spleens included CD19+ B cells (including IgD+ unswitched naïve B cells), NK1.1+ NK cells, and CD3+ T cells (Figure 2C). Total T cell populations as a percentage of CD45+ cells decreased in Cxcr2 KO spleens, including CD8+ cytotoxic T cells and CD4+ helper T cells. In contrast, immunosuppressive FoxP3+ CD25+ regulatory T cells and NKT cells (CD3+ NK1.1+) increased as percentage of total CD45+ cells, consistent with impaired adaptive immune homeostasis (Figure 2D). Within the CD3+ T cell population, T cell subsets differed in Cxcr2 WT and KO spleen. NKT cells increased, while cytotoxic (CD8+) and helper (CD4+) T cells decreased, and T regs (FoxP3+ CD25+) were unchanged (Figure 2F).

Together, these splenic changes suggest a shift toward an immunosuppressive, innate-skewed immune environment, including neutrophils, MDSCs, and M2 macrophages, with increased regulatory T cells and NKT cells, at the expense of adaptive effector populations, such as CD8+ T cells and B cells, a pattern that is further reflected in the bone marrow.

In the bone marrow, the myeloid compartment significantly expanded in Cxcr2 KO mice compared to WT mice, from ∼84% of CD45+ cells in WT and nearly 98% in Cxcr2 KO bone marrow (Figure 2C). This increase in total myeloid cells arose largely from granulocytic neutrophils and PMN-MDSCs. In contrast, M-MDSCs, total macrophages, and M2 macrophages decreased in KO bone marrow (Figure 2D). Within the macrophage compartment, we saw a corresponding polarization shift toward an M1-like marker expression (Figure 2E).

The lymphoid populations in the bone marrow paralleled those seen in the spleen (Figure 2C). Similarly, B cells markedly decreased, including naïve/unswitched B cell populations. NK cells and T cells decreased globally, including CD3+ T cells, CD8+ cytotoxic T cells, CD4+ helper T cells, and FoxP3+ regulatory T cells (Figure 2F). In contrast, NKT cells were unchanged as a proportion of CD45+ cells and significantly increased as a proportion of total CD3+ cells in Cxcr2 KO bone marrow (Figure 2F).

Together, these data suggest that Cxcr2 deficiency skews the bone marrow toward myelopoiesis with suppression of adaptive immune populations. The net effect of Cxcr2 deficiency across both tissues is an expansion of suppressive and regulatory cells and loss of adaptive immune effector populations, suggesting a systemic shift toward immune tolerance rather than cytotoxic surveillance.

### Mammary gland branching morphogenesis is delayed in Cxcr2 KO mice throughout development

To examine the effects of Cxcr2 deficiency on mammary gland development, we next evaluated branching morphogenesis in carmine-stained mammary gland whole mounts (Figure 3A). We quantified the distance of ductal invasion into the mammary fat pad as well as the total mammary epithelial area in 3-, 5-, 8-, and 12-week-old mice (Figure 3B-C).

**Fig. 3.**
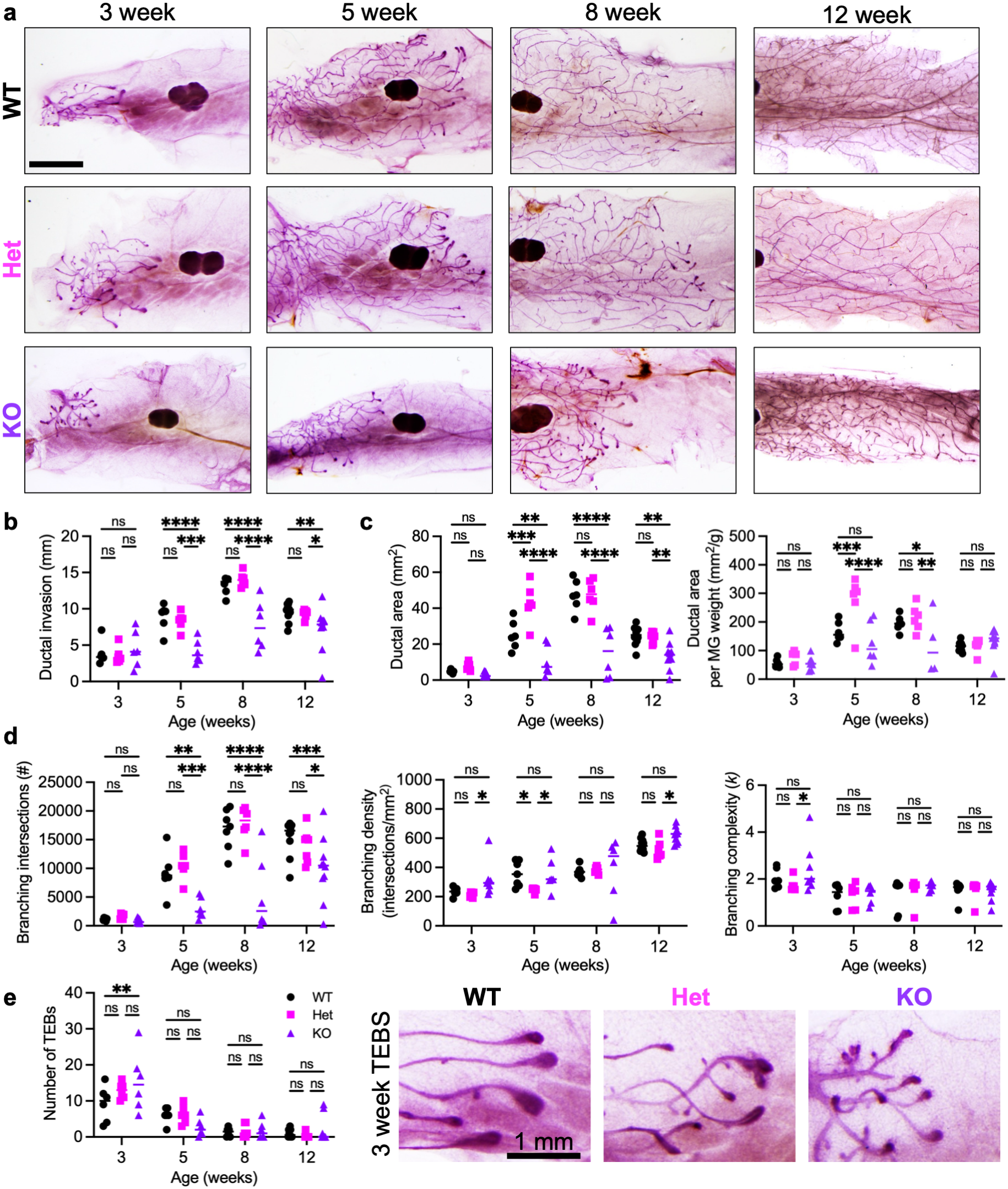
Cxcr2 KO mice display delayed ductal outgrowth and reduced branching during pubertal mammary development. (A) Representative carmine-stained whole mounts of the fourth abdominal mammary gland from Cxcr2 WT, Het, and KO mice at 3, 5, 8, and 12 weeks; scale bar = 3 mm. (B) Ductal invasion (mm), (C) ductal area (mm^2^), and (D) Sholl analysis of branching intersections (#), branching density (intersections/mm^2^) and branching complexity (k) at each age. (E) Number of terminal end buds (TEBs) per gland at each age (left) and representative images of 3-week TEBs from each genotype (right; scale bar = 1 mm). n = 6-9/genotype/age. Statistical analysis by two-way ANOVA with Tukey’s multiple comparisons test. Lines indicate mean. ns, not significant; *p < 0.05, **p < 0.01, ***p < 0.001, ****p < 0.0001

Mammary epithelial branching was significantly delayed with age in Cxcr2 KO mice. In 3-week mice, ductal invasion did not significantly differ by genotype, and the rudimentary ductal epithelium appeared grossly normal. At 5 weeks, the average ductal invasion in Cxcr2 KO was 4.09 mm versus 8.88 mm for WT. At 8 weeks old, the average ductal invasion distance was 7.74 mm versus 13.20 mm in Cxcr2 KO and WT mice, respectively. Thus, by 8 weeks of age, the distance of the mammary epithelial invasions decreased by 41% in Cxcr2 KO mice (P<0.0001).

Similarly, the total area of the branching epithelium decreased in the Cxcr2 KO mice compared to WT mice as they aged (P<0.0001). While not significant at 3 weeks old, by 5 weeks of age, Cxcr2 KO ductal area was 10.77 mm^2^ on average, versus 25.12 mm^2^ in WT, which is a 57% reduction in ductal area. By 8 weeks of age, the ductal area decreased 67% in Cxcr2 KO mammary glands versus WT (15.42 mm^2^ versus 47.25 mm^2^, respectively).

To quantify the ductal branching, we used a Sholl analysis plugin in ImageJ to create maps of branching epithelium in whole mounts to determine the total number of branching intersections, branching density, and branching complexity (Figure 3D) [22]. The number of branching intersections decreased in the 5- and 8-week Cxcr2 KO mice, congruent with reduced epithelial area and ductal invasion. However, the branching density and complexity (rate of branching as a function of distance from nipple) were not consistently altered with Cxcr2 deletion.

Manual counting of terminal end buds (TEBs) revealed a moderate increase in TEB number in 3-week Cxcr2 KO and Het glands versus WT, while no differences in TEBs were seen in 5-, 8-, and 12-week-old glands (Figure 3C). TEB width also did not significantly differ with Cxcr2 status (Supplemental Figure 3A).

### Transplantation studies with Cxcr2 WT and KO epithelium do not recapitulate the defects in naïve Cxcr2 KO mice

We next wanted to assess if the decreased mammary gland branching morphogenesis in Cxcr2 KO mice was due to defects in the mammary epithelium, dependent on the stroma, or both. We transplanted primary mammary epithelial cells (MECs) from WT and Cxcr2 KO donor mice into the cleared mammary fat pads of Cxcr2 WT and KO recipient mice. Mammary glands were collected 5 weeks after transplantation, based on the timepoint where a significant difference in the branching epithelium was evident in Cxcr2 KO mice (Figure 4A). The tissues were carmine stained for whole mount analysis, and the glands were analyzed for mammary gland branching. Glands with no epithelial growth were excluded from analysis as failed injections.

**Fig. 4.**
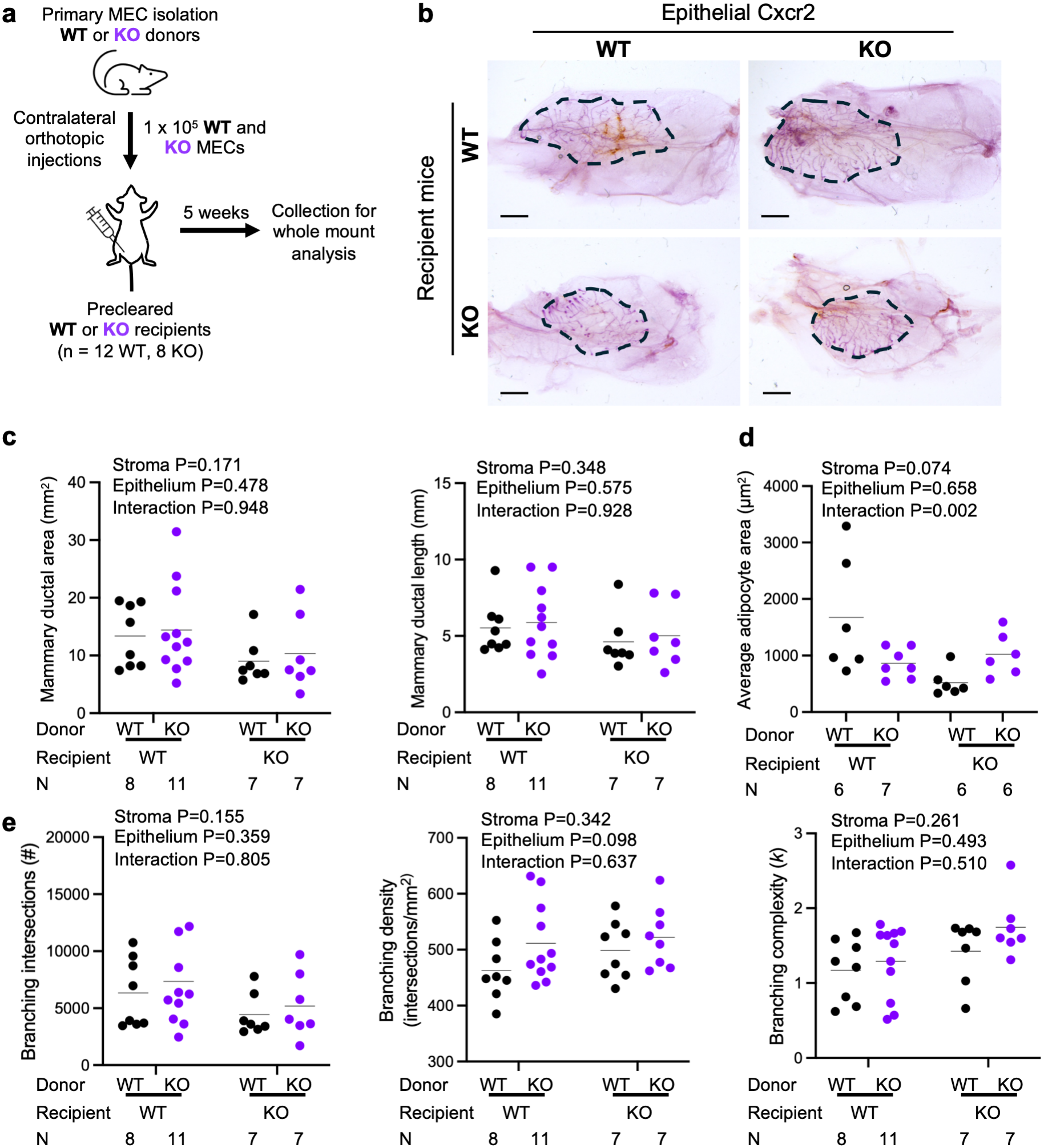
Transplanted Cxcr2 status does not recapitulate the branching phenotype of the naïve KO host. (A) Schematic of the transplant design: primary mammary epithelial cells (MECs; 1 x 10^5^) from Cxcr2 WT or KO donors were injected into contralateral precleared fat pads of Cxcr2 WT or KO recipients (n = 12 WT, 8 KO recipients) with whole-mount analysis 5 weeks post-transplant. (B) Representative carmine-alum whole mounts of transplanted glands across donor/recipient genotype combinations; scale bar = 3 mm. (C) Mammary ductal area (mm^2^, left) and ductal length (mm, right). (D) Average adipocyte area (µm^2^). (E) Sholl analysis of branching intersections (#), branching density (intersections/mm^2^), and branching complexity (k). Ductal area, length, density, and complexity were analyzed by linear mixed-effects models, while total branching intersection counts were log-transformed and analyzed using a Gaussian linear mixed-effects model. All models included stromal (recipient) and epithelial (donor) Cxcr2 genotype and their interaction as fixed effects, with a random effect for recipient mouse to account for paired samples. P-values for the stroma, epithelium, and interaction terms are shown on each panel. N per donor/recipient combination is indicated below each graph. Lines indicate mean

Quantification of the epithelial area from whole mounts did not differ based on Cxcr2 WT or KO status from either the epithelial donor cells or recipient stroma (Figure 4B-C). Because the transplant cohorts included paired and unpaired samples across studies, mammary ductal area was analyzed using a linear mixed-effects model with stromal (recipient) and epithelial (donor) Cxcr2 genotype as fixed effects. Mammary ductal area was not significantly impacted by stromal Cxcr2 status (P=0.171), epithelial Cxcr2 status (P=0.478), or their interaction (P=0.948). In Cxcr2 WT recipient mice, the mammary ductal area was 13.40±5.41 mm^2^ with WT MECs and 14.41±7.89 mm^2^ with KO MECs. In Cxcr2 KO recipient mice, the mammary ductal area was 9.02±3.92 mm^2^ with WT MECs and 10.36±6.49 mm^2^ with KO MECs. Thus, neither epithelial nor stromal Cxcr2 status significantly impacted mammary ductal area, and WT:WT and KO:KO combinations did not differ despite the marked branching area defect observed in the naïve KO host. Mammary ductal invasion (Figure 4C) also did not differ based on stromal (P=0.348) or epithelial (P=0.575) Cxcr2 status (interaction P=0.928).

We next quantified adipocyte area in the mammary fat pad of transplant recipient mice. Neither recipient genotype (P=0.074) nor donor genotype (P=0.658) had a significant effect on adipocyte area. However, there was a significant impact of their interaction (P=0.002; Type III test), indicating that the effect of epithelial Cxcr2 genotype depended on the stromal genotype of the recipient. In WT recipients, KO donor epithelium was associated with 42% lower adipocyte area than WT donor epithelium, whereas in KO recipients, KO donor epithelium was associated with 98% higher adipocyte area (Figure 4D), indicating that adipocyte size depends on the combination of epithelial and stromal Cxcr2 status.

We also measured mammary epithelial branching intersections, density, and complexity by Sholl analysis (Figure 4E). All three outcomes were analyzed using linear mixed-effects models, with total branching intersections log-transformed and the other two outcomes analyzed on their original scales. Neither stromal Cxcr2 status (P=0.155), epithelial Cxcr2 status (P=0.359), nor their interaction (P=0.805) reached statistical significance for branching intersections. Branching density (stroma P=0.342, epithelium P=0.098, interaction P=0.637) and branching complexity (stroma P=0.261, epithelium P=0.493, interaction P=0.510) also did not differ significantly based on stromal or epithelial Cxcr2 status.

Together, the lack of statistically significant genotype effects on ductal area, invasion length, or branching intersections, density, and complexity, combined with the significant epithelial-stromal interaction for adipocyte size, indicates that transplantation does not recapitulate the reduced branching of the naïve KO host. These findings suggest that the role of Cxcr2 in supporting mammary epithelial growth may depend on developmental timing or other factors, such as immune responses, extracellular matrix, or vasculature.

### Cxcr2 deletion produces smaller adipocytes in the mammary fat pad

The mammary fat pad is composed primarily of adipocytes and collagen-rich extracellular matrix that crosstalk with the invading mammary epithelium. To assess if Cxcr2 deletion impacts the organization of the mammary fat pad, we employed histological analysis. H&E staining of mammary glands revealed a notable reduction in adipocyte size in the mammary fat pad of Cxcr2 KO mice (Figure 5A). We quantified adipocyte number per field of view and average adipocyte size. Adipocytes were significantly smaller by adipocyte area in Cxcr2 KO mice at all ages, with a corresponding increase in number per field, compared to WT (Figure 5B). Cxcr2 KO adipocytes also decreased in average area as they aged, in contrast to adipocytes from WT littermates, which did not significantly change in size. Specifically, the mean adipocyte area in 3-week-old Cxcr2 KO mammary glands was only 1313.47 μm^2^, compared to 1849.68 μm^2^ in 3-week-old WT mice. At 5 weeks, the Cxcr2 KO average adipocyte area was 933.18 μm^2^ versus 1902.70 μm^2^ for WT. At 8 weeks, the average adipocyte area was 595.91 μm^2^ versus 2088.92 μm^2^ in Cxcr2 KO and WT mice, respectively. While KO adipocyte area was 71% of WT in 3-week animals, by 8 weeks of age, Cxcr2 KO mammary adipocytes were only 29% of the average area of WT controls (P<0.0001).

**Fig. 5.**
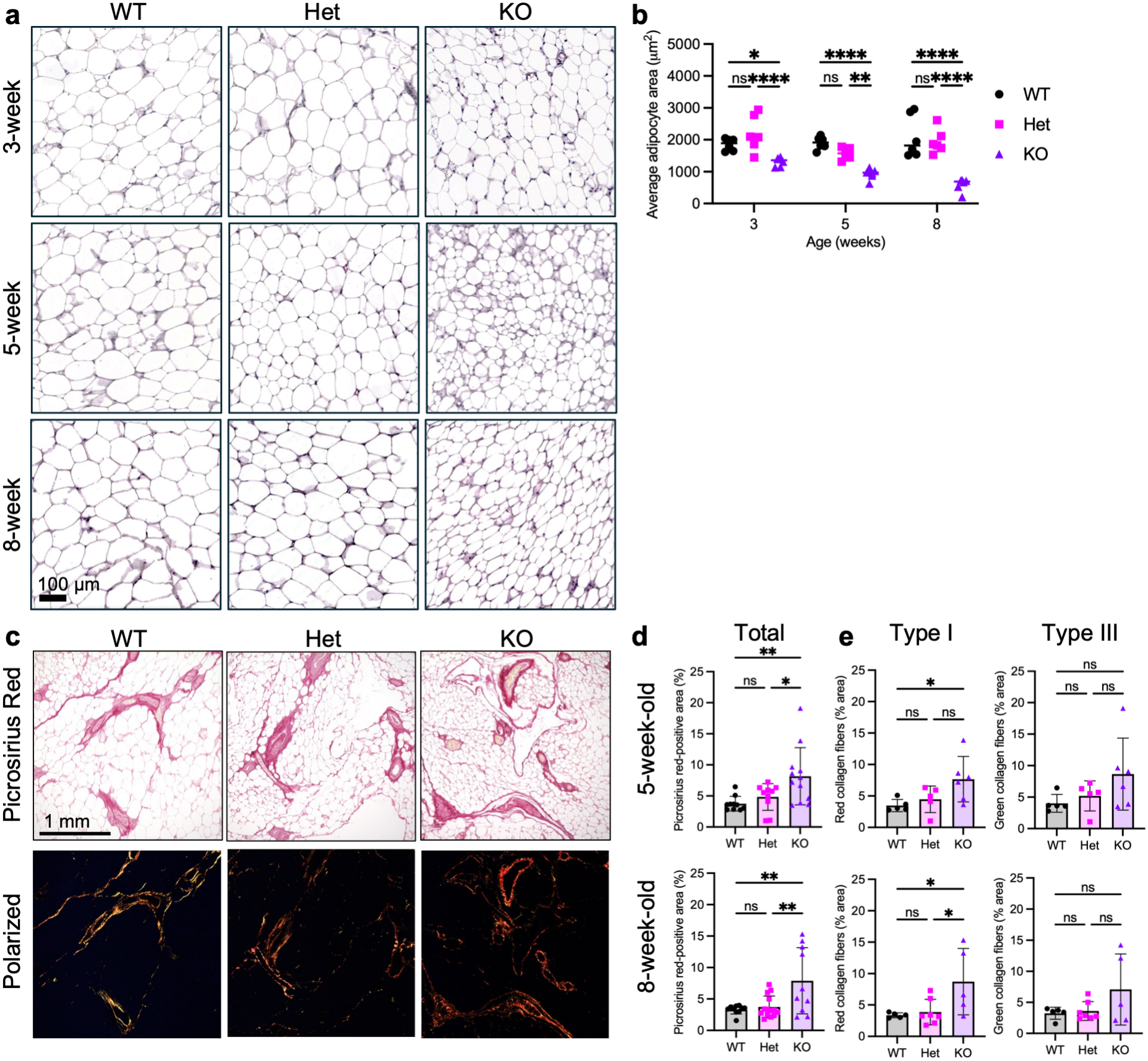
Cxcr2 deletion produces smaller adipocytes and increased collagen fiber content in the mammary fat pad. (A) Representative H&E images of mammary fat pad from Cxcr2 WT, Het, and KO mice at 3, 5, and 8 weeks; scale bar = 100 µm. (B) Average adipocyte area (µm^2^) at each age (n = 6/genotype/age). (C) Representative picrosirius red–stained sections imaged by brightfield (top) and polarized light (bottom) in 8-week Cxcr2 WT, Het, and KO glands; scale bar = 1 mm. (D) Quantification of total picrosirius red-positive area (%) and (E) type I (red birefringent) and type III (green birefringent) collagen fibers (% area) in 5-week-old (top) and 8-week-old (bottom) glands (n = 5-7/genotype). Statistical analyses by ANOVA with Tukey’s multiple comparisons test. Lines indicate mean. ns, not significant; *p < 0.05, **p < 0.01, ***p < 0.001, ****p < 0.0001

### Cxcr2 deletion increases collagen in the mammary stroma

We quantified the collagen content in 5- and 8-week-old Cxcr2 WT, Het, and KO mammary glands via picrosirius red staining (Figure 5C). Sections were then imaged under brightfield and polarized light to quantify the birefringent collagen content per mammary tissue area. Type I collagen (thick, rigid fibers) and type III collagen (thin, flexible fibers) were identified by red and green birefringence, respectively [23, 24]. Interestingly, Cxcr2 KO collagen fiber content was significantly elevated when normalized to the total analyzed mammary gland tissue area (Figure 5D). In 5-week-old mice, the total picrosirius-red stained area increased significantly in Cxcr2 KO (P=0.0067), elevated from a mean percentage of 3.481% in WT and 4.478% in Het to 7.664% in Cxcr2 KO mammary glands. Red collagen fiber area significantly increased (P=0.0449), while the increase in green collagen fiber was variable in Cxcr2 KO samples (Figure 5E). In 8-week-old mice, the mean total picrosirius-red stained area was 3.305% in WT and 3.737% in Het compared to 7.898% in KO (P=0.0060, one-way ANOVA; Figure 5D). As in 5-week-old mice, red birefringence from thick type I collagen fibers significantly increased in Cxcr2 KO glands in 8-week-old mice (P=0.0281), while green fibers varied (Figure 5E). This indicates that Cxcr2 KO mammary glands have increased bulk collagen content, and particularly thick type I-like collagen fibers.

### Mammary epithelial organization, apoptosis, and proliferation are preserved in Cxcr2 KO mice

To examine if the organization of the mammary epithelium is altered in Cxcr2 KO mice, we assessed the luminal marker keratin-8 (K8) and myoepithelial markers keratin-14 (K14) and smooth muscle actin (SMA) by immunofluorescence imaging. As Cxcr2 Het mice were phenotypically indistinguishable from WT throughout this study, only Cxcr2 WT and KO mammary glands were analyzed by immunofluorescence staining. We co-stained for K8 and SMA (Figure 6A) or K8 and K14 (Figure 6B-C) in 8-week-old mammary gland tissue. Analysis revealed no detectable differences in the organization of the Cxcr2 KO epithelial layers compared to WT mice. Quantification of K8+, K14+, and double-positive (K8+K14+) cells (as a percentage of total epithelial cells per duct), the percentage of ductal area occupied by each marker, and mean fluorescence intensity per ductal area similarly did not differ significantly between Cxcr2 WT and KO ducts (Figure 6C).

**Fig. 6.**
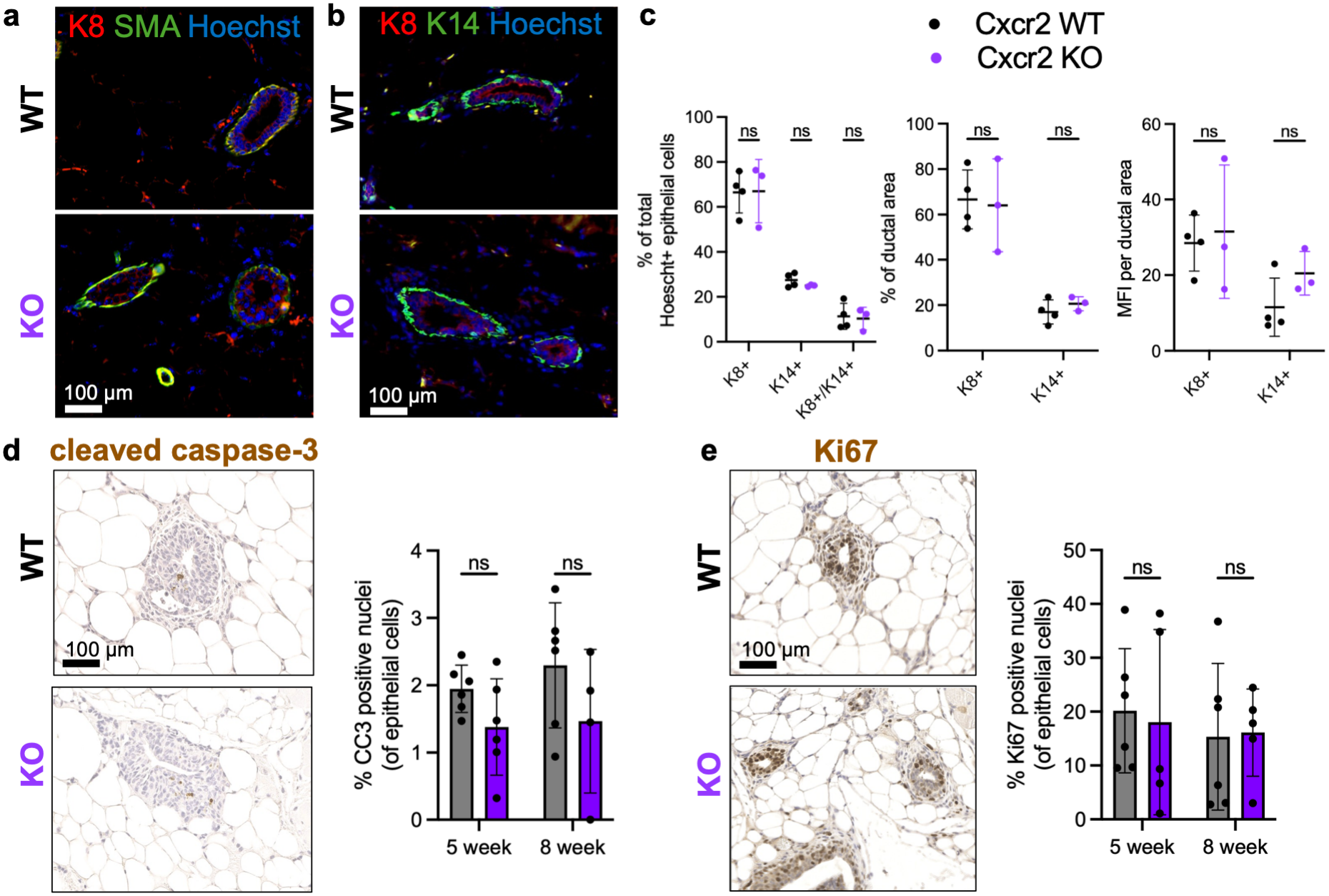
Mammary epithelial architecture, proliferation, and apoptosis are preserved in Cxcr2 KO mice. (A) Representative immunofluorescence of 8-week-old Cxcr2 WT and KO mammary ducts stained for keratin-8 (K8, red), smooth muscle actin (SMA, green), and nuclei (Hoechst, blue); scale bar = 100 µm. (B) Representative immunofluorescence stained for K8 (red), keratin-14 (K14, green), and nuclei (Hoechst, blue); scale bar = 100 µm. (C) Quantification of K8+, K14+, and K8+K14+ double-positive cells (as % of total Hoechst+ epithelial cells), the percentage of ductal area occupied by K8+ or K14+ signal, and mean fluorescence intensity (MFI) per ductal area, in 5-week-old Cxcr2 WT and KO ducts (n = 4 WT, 3 KO). (D) Representative cleaved caspase-3 (CC3) immunohistochemistry in Cxcr2 WT, Het, and KO glands; scale bar = 100 µm. (E) Quantification of CC3+ nuclei (as % of epithelial cells) at 5 weeks (left) and 8 weeks (right) (n = 4-6/genotype). Statistical analysis by ANOVA with Tukey’s multiple comparisons test. (F) Representative Ki67 immunohistochemistry in 5-week-old Cxcr2 WT and KO glands; scale bar = 100 µm. (G) Quantification of Ki67+ nuclei (as % of epithelial cells) at 5 weeks (left) and 8 weeks (right) (n = 4-6/genotype). Statistical analysis by Welch’s t-test. Lines indicate mean. ns, not significant

These findings indicate that epithelial lineage composition and ductal architecture, as assessed by keratin-8, keratin-14, and smooth muscle actin staining, were not detectably altered by Cxcr2 loss, suggesting that Cxcr2 is not required for lineage specification or epithelial identity, but rather affects growth dynamics within an otherwise normally organized epithelium.

To determine if Cxcr2 deletion impacts cell death or proliferation of the developing mammary epithelium, we employed IHC staining for cleaved-caspase 3 (CC3) or Ki67, respectively (Figure 6D-G). Staining for cleaved caspase-3 revealed no changes in apoptosis levels based on Cxcr2 status, with a mean percentage of 1.94±0.35% CC3+ cells in WT, 2.92±1.84% in Het, and 1.38±0.72% in KO glands of 5-week-old mice (P=0.0983), and 2.30±0.93% CC3+ cells in WT, 2.15±0.39% in Het, and 1.47±1.06% in KO mammary epithelium of 8-week-old mice (P=0.2888, ANOVA; Figure 6D-E). The proportion of Ki67+ epithelial cells did not differ between Cxcr2 WT and KO mammary ducts at either 5 or 8 weeks of age (Figure 6F-G; P=0.4286 and P=0.5273, respectively; n = 4-6/genotype), indicating that loss of Cxcr2 does not detectably alter intrinsic mammary epithelial proliferation.

## Discussion

In this study, we characterized the role of Cxcr2 in postnatal mammary gland development using a germline Cxcr2 knockout model on the FVB/N background. We found that Cxcr2 KO mice exhibited delayed ductal outgrowth and reduced epithelial area during puberty, accompanied by pronounced stromal remodeling—smaller adipocytes and increased collagen deposition—and severe systemic immune dysregulation marked by expansion of granulocytic and suppressive myeloid populations and loss of adaptive immune cells. Despite these defects, mammary epithelial lineage organization, proliferation, and apoptosis were preserved, and branching complexity was largely intact. Transplantation of Cxcr2 KO epithelium or stroma did not recapitulate the branching defect of the intact KO host. Together, these findings indicate that Cxcr2 regulates mammary ductal outgrowth not through an epithelial cell-intrinsic mechanism, but by shaping the immune and stromal microenvironment in which the epithelium develops.

A defining feature of the Cxcr2 KO FVB/N phenotype was global impairment of mouse development, including decreased body size, decreased adiposity, and increased mortality. Cxcr2 KO mice on this background also exhibit significant skeletal defects, including loss of trabecular and cortical bone and increased osteoclast-mediated resorption that we characterized previously [21]. These systemic defects complicate the interpretation of mammary-specific phenotypes but are nonetheless biologically informative because they underscore the importance of CXCR2 signaling in coordinating organismal growth, immune homeostasis, and tissue development. Importantly, mammary gland defects in Cxcr2 KO mice exceeded what would be predicted based on reduced body weight alone, indicating a specific requirement for CXCR2 in mammary gland expansion beyond generalized growth delay.

Whole mount and quantitative morphometric analyses revealed a marked delay in mammary branching morphogenesis in Cxcr2 KO mice, characterized by reduced ductal invasion and epithelial area. While the absolute number of branching intersections decreased with Cxcr2 deficiency, branching density and complexity were largely preserved, suggesting that the primary defect lies in ductal outgrowth and tissue expansion, rather than in intrinsic branching patterning. The preservation of branching complexity, despite severely reduced total area, implies that the epithelial branching program itself is intact in Cxcr2 KO mice and that the limiting factor may be stromal permissiveness for ductal invasion, consistent with the observed adipocyte and collagen defects, rather than a failure of epithelial patterning. The transient increase in terminal end buds observed at early timepoints may reflect a delayed developmental progression or an impaired transition from early invasion to mature ductal architecture.

Transplantation experiments further support a collaborative role for epithelial and stromal Cxcr2 in mammary development and support a role for Cxcr2 in establishing the microenvironmental niche that supports epithelial growth. Neither epithelial genotype nor stromal genotype alone was sufficient to recapitulate the branching defects observed in naïve Cxcr2 KO mice, consistent with the interpretation that mammary phenotypes arise from a disrupted niche that is present only in the intact developing mammary gland or from systemic or multicompartmental defects inherent to the germline knockout. This suggests that Cxcr2 contributions to mammary gland development are complex, are not driven exclusively by either the epithelium or the stroma, and are driven by factors that differ or are disrupted between normal development and transplantation. The experimental parameters of transplantations are known to induce a wound healing and immune response, reorganize the extracellular matrix, and disrupt the local vasculature, all of which could differ from the context during normal mammary development. While technical limitations due to the mortality and fragility of the Cxcr2 mice restricted the sample number of our transplantation studies into the Cxcr2 KO stroma, these data align with the broader conclusion that CXCR2 signaling in stromal compartments indirectly shapes epithelial morphogenesis. Future transplantation experiments could overcome the limiting comorbidities of the Cxcr2 KO host by collecting, mixing, and transplanting the isolated epithelium and mammary fat pad into the kidney capsule of Cxcr2 WT and KO mice to determine the specific impacts of the epithelium and stroma during pubertal development. In addition, experiments using bone marrow transplantation experiments that swap the marrow of WT and KO mice will determine if the immune system or other stroma cells are essential in the Cxcr2-mediated developmental response.

A striking feature of Cxcr2 KO mice was their severe immune dysregulation, including expansion of granulocytic and suppressive myeloid populations and loss of adaptive immune cells in both spleen and bone marrow. This immune imbalance mirrors phenotypes of Cxcr2 deficiency described in other tissues and disease contexts [25]. This imbalance is particularly relevant in the mammary gland, where macrophages, neutrophils, and other immune cells play essential roles in terminal end bud formation, ductal elongation, and extracellular matrix remodeling. Disruption of immune cell recruitment, polarization, or function could contribute directly to impaired branching morphogenesis in Cxcr2 KO glands. Macrophage depletion studies established that ductal elongation requires macrophage-TEB interactions [3, 4, 26]. Our observed shift toward M2-polarized macrophages in the Cxcr2 KO spleen and the reduction in macrophages and M-MDSCs in KO bone marrow suggest that macrophage function and trafficking are specifically perturbed. Whether the macrophage population within the KO mammary gland fat pad is similarly dysregulated, and whether restoration of normal macrophage balance could rescue ductal elongation, are important questions for future work. Notably, immune profiling of the mammary gland itself was not performed in this study, representing an important future direction to determine whether systemic immune alterations translate into local stromal defects during pubertal development.

The contribution of the CXCR2-dependent mammary stroma to local epithelial development has not been examined previously except in one study that evaluated the mammary glands of 13-week-old mice and the ability of tissue from mammary glands and ovaries of Cxcr2 KO mice to be transplanted into immune competent mice [20]. In that study, the Cxcr2 KO epithelium appeared smaller at 13 weeks than in WT, but ductal invasion, other branching parameters, and earlier developmental time points were not quantified. In addition, Cxcr2 KO epithelium transplanted into the WT mammary fat pad in the BALB/c background gave rise to a healthy, fully branched mammary gland tissue that the team attributed to inflammation-induced alterations to pituitary hormone and progesterone signaling in the Cxcr2 KO mouse host. Because our animal facility is barrier-maintained under only specific pathogen-free conditions and is not a complete barrier facility, we cannot rule out environmental factors in our experiments.

The immune and skeletal phenotypes that we have observed in Cxcr2 KO mice may reflect a common disruption across progenitor lineages. Osteoclasts derive from the same monocyte/macrophage precursor pool as tissue macrophages. The elevated osteoclast activity and increased bone resorption observed in Cxcr2 KO FVB/N mice [21] are consistent with the myeloid expansion and skewed macrophage polarization identified in this study, suggesting that Cxcr2 coordinates myeloid cell fate across tissue compartments. In parallel, the reduced mammary and impaired bone homeostasis in Cxcr2 KO mice raises the possibility of a broader mesenchymal linage defect, since Cxcr2 signaling supports adipogenic differentiation from MSC precursors [18, 19]. Whether Cxcr2 deficiency shifts MSC fate between adipogenic and osteogenic lineages in a tissue-dependent manner remains an open question but would suggest a role for Cxcr2 in coordinating mesenchymal stromal homeostasis across tissues.

In addition to immune alterations, Cxcr2 KO mammary glands exhibited additional pronounced stromal defects, including reduced adipocyte size and increased collagen deposition. Mammary adipocytes provide both structural support and paracrine signals that facilitate ductal invasion [27]. The progressive reduction in adipocyte size with age in Cxcr2 KO mice, coupled with elevated collagen fiber content, suggests a stiffer, less permissive stromal environment that may mechanically and biochemically constrain epithelial outgrowth. Altered type I collagen deposition is associated with impaired ductal elongation and altered epithelial-stromal crosstalk in mouse mammary gland [28–30] and in human breast cancers [31–33]. The increase in thick, red-birefringent (type I-like) collagen fibers is particularly notable because type I collagen increases stromal stiffness and regulates epithelial cell migration and invasion through integrin-mediated mechanosensing pathways [30, 34, 35]. Stromal fibroblasts are the primary cellular source of collagen in the mammary gland, and TGF-β signaling is a well-established driver of fibroblast activation and collagen synthesis [36]. CXCR2-dependent regulation of fibroblast activity or TGF-β signaling may therefore contribute to the excess collagen deposition observed in Cxcr2 KO mammary glands, since both pathways are known downstream targets of CXCR2 in stromal remodeling contexts [37, 38]. These findings indicate that Cxcr2 loss disrupts the balance between adipogenesis and extracellular matrix remodeling, further reinforcing the conclusion that mammary epithelial defects arise indirectly from stromal dysfunction.

A limitation of this study is the reliance on a germline Cxcr2 knockout model, which introduces developmental compensation, systemic immune dysregulation, and endocrine abnormalities that complicate attribution of mammary phenotypes to specific cell types. Mouse strain-specific differences also impact the immune signaling and thus the interpretation of specific CXCR2 functions in the developing mammary gland. Previous studies have demonstrated pituitary dysfunction and altered steroid hormone signaling in Cxcr2 KO mice [20], raising the possibility that endocrine defects contribute to impaired mammary development in this model. Hormone levels were not measured in this study, and our facility pathogen testing showed the presence of pathogen species that may alter Cxcr2 KO phenotypes [20, 39]; thus, endocrine contributions due to chronic inflammation in the Cxcr2 KO mice cannot be excluded. Additionally, the poor health and increased mortality of Cxcr2 KO mice limited the feasibility of transplantation experiments and sample sizes. These technical limitations underscore the need for conditional, tissue-specific deletion strategies to disentangle epithelial, stromal, immune, and endocrine contributions to mammary gland development. Targeted lineage-specific deletions of Cxcr2 have been assessed in myeloid cells during tumor progression [40] and in oligodendrocytes in the developing spinal cord [41], which enabled the interpretation of CXCR2 signaling without the confounding developmental defects of the germline KO mouse. Future studies employing immune cell-specific or stromal-specific deletion of Cxcr2, coupled with direct profiling of mammary-resident immune populations, will be critical for defining the precise mechanisms by which CXCR2 regulates pubertal mammary development.

Collectively, these findings position CXCR2 as a developmental regulator of tissue remodeling crosstalk and immune balance, rather than as a direct epithelial growth factor. These observations also carry implications that extend beyond normal development. The stromal and immune features remodeled by Cxcr2 loss during puberty, including expansion of granulocytic and myeloid-derived suppressor cell populations, M2-skewed macrophage polarization, suppression of adaptive effector cells, and accumulation of thick type I collagen fibers, are each recognized hallmarks of the pro-tumorigenic breast microenvironment, where CXCR2 signaling promotes tumor growth, metastasis, and immunosuppression. Our data therefore suggest that the same CXCR2-dependent programs that orchestrate immune balance and stromal architecture during normal ductal morphogenesis may be co-opted in the breast tumor microenvironment, where dysregulated chemokine signaling reshapes the stroma to favor progression. Defining whether CXCR2 acts through shared immune and stromal intermediaries in the developmental and malignant settings will be an important direction for future work and may clarify how a receptor essential for normal mammary patterning contributes to tumor-associated remodeling.

## Methods

### Animals

Cxcr2 KO (B6.129S2(C)-*Cxcr2*^tm1Mwm^/J) mice were acquired from Jackson Laboratories (JAX: 006848) and backcrossed at least 10 generations to the FVB/N background (Charles River, 559NCIFVB) [21].

Mice were barrier-maintained under specific pathogen-free conditions in the University of Notre Dame Freimann Life Sciences animal facility. All animal studies were conducted with approval from the University of Notre Dame Institutional Animal Care and Use Committee (protocols 21-11-6912, 21-12-6910, 21-12-6936, 21-12-6937, 24-10-8880, 24-10-8883, 24-10-8884) and in accordance with guidelines from the National Institutes of Health and U.S Department of Defense.

### Pathogen testing

Fresh fecal samples from Cxcr2 WT and KO mice (n=2/genotype) were submitted to IDEXX BioAnalytics for pathogen monitoring using the Opti-XXpress/EDx Mouse Prevalent Profile. Pathogens evaluated by PCR included: Aspiculuris tetraptera, EDIM, Entamoeba muris, Helicobacter bilis, Helicobacter ganmani, Helicobacter hepaticus, Helicobacter mastomyrinus, Helicobacter rodentium, Helicobacter spp., Helicobacter typhlonius, MHV, MNV, MPV, MVM, Myocoptes, Radfordia/Myobia, Rodentibacter heylii, Rodentibacter pneumotropicus, Spironucleus muris, Syphacia muris, Syphacia obvelata, TMEV.

### Body composition analysis

Whole body composition was measured in 3-, 5-, 8-, and 12-week-old female Cxcr2 WT, Het, and KO mice with whole-body quantitative magnetic resonance using an EchoMRI Body Composition Analyzer (EchoMRI LLC, Houston, TX).

### Mammary fat pad preclearing

For transplantation experiments, endogenous mammary epithelium was surgically cleared from the right and left fourth mammary fat pads of 3-week-old recipient Cxcr2 WT and KO FVB mice. Incisions were closed with Relex 7 mm wound clips (Stoelting Co., Cat. #59034) using a wound clip stapler (Stoelting, Cat. #59038).

### Isolation of primary mammary epithelial cells

Primary mammary epithelial cells (MECs) were isolated from the numbers 2/3 and 4 mammary glands of 8–16-week-old donor Cxcr2 WT and KO mice by differential centrifugation as used previously [42–44]. Mice were euthanized by CO_2_ inhalation, and mammary glands were dissected with muscle and lymph node removed. Tissues were weighed, pooled by genotype, and placed in a 15 mL tube containing 3 mL wash buffer (10 mM HEPES, 2.5% FBS in RPMI 1640 (Gibco, Cat. #61870). In a culture hood, the glands were chopped with #10 scalpels for 2-5 minutes and transferred to fresh tubes with collagenase buffer (2 mg/mL Collagenase IV (Sigma, Cat. #C5138) in wash buffer; 5 mL buffer per gram of tissue). Tissues were incubated at 37°C with moderate shaking (200 RPM; New Brunswick Innova 2300/2500, Eppendorf) for 1-1.5 hours until the solution appeared homogenous without large clumps. Organoids were pelleted by centrifugation at 600 x g for 5 minutes and the fatty supernatant was aspirated. Organoids were further enriched by 4-6 pulse-spin washes in collagenase buffer (pulsed to 500 x g with maximum deceleration), continued until red blood cells were absent from the pellet and ≥90% of the preparation consisted of organoids. The organoid pellet was then resuspended in 2 mL of 0.5 g/L trypsin/EDTA in a well of a 6-well plate and incubated at 37°C for 15–20 minutes with occasional pipetting and microscopic monitoring. Eight mL of wash media was added, and the suspension was pelleted at 450 x g for 3 minutes. The pellet was resuspended in 5-10 mL serum-free DMEM/F12 containing DNase (100 μg/mL final concentration) and mixed by slow inversion for 2-3 minutes at room temperature. Cells were pelleted at 450 x g for 3 minutes, resuspended in 10 mL wash media, and filtered through a 70 μm cell strainer (STEMCELL Technologies, Cat. #27216) to remove aggregates. The wash was repeated, and cells were resuspended in 5 mL growth media and counted by hemocytometer. Growth media consisted of DMEM/F12 supplemented with 5 μg/mL insulin, 1 μg/mL hydrocortisone, 10 ng/mL EGF (Gibco, Cat. #PMG8041), 10% FBS, 100 U/mL Pen/Strep, and 50 μg/mL Gentamicin. Cells were maintained in non-adherent culture at 37°C and used the same day for orthotopic transplantation.

### Orthotopic mammary transplantation

Primary MECs were counted by hemocytometer, resuspended at 1×10^5^ cells per 20 μL DPBS with Ca^2+^ and Mg^2+^ (Sigma-Aldrich, Cat. #D5652-10X1L), and kept on ice prior to injection. Cells were injected contralaterally into the precleared mammary fat pads of recipient mice as described previously [45]. Animals were sacrificed 5 weeks post-transplantation and mammary glands were collected for whole mount analysis. In this experiment, the injection success rate limited the number of paired samples that were included for analysis.

### Flow cytometry

Spleens were harvested into RPMI supplemented with 10% FBS and dissociated by mechanical digestion. Tissue was crushed in a cell culture dish with the back of a 5 mL syringe plunger in staining media (PBS + 0.2% BSA + 2% FBS + 1 mM EDTA) and passed through a 70 μm cell strainer. The remaining tissue was further washed through the 70 μm filter with the black, rubber end of the plunger using staining media. The filtered suspension was centrifuged at 500xg for 5 minutes at 4°C, and red blood cells were lysed by resuspending the pellet in 3 mL eBiosciences RBC lysis (00-4300-54) buffer for 5 minutes on ice. Lysis was quenched with 20 mL cold staining media, and cells were passed through a 40 μm strainer, centrifuged again, and resuspended in 1-2 mL staining media. Bone marrow was isolated from femurs and tibiae under sterile conditions. Marrow was flushed with 5 mL complete RPMI (RPMI 1640 + 10% FBS + 2 mM L-glutamine) using a 25g needle and syringe, passed through a 70 μm strainer by mechanical dissociation, and centrifuged at 500xg for 4 minutes at 4°C. Red blood cells were lysed as described above, quenched with 15 mL cold staining media, filtered through a 40 μm strainer, and resuspended in 1-2 mL staining media.

For staining, 1×10^6^ cells per well were plated in a 96-well plate and washed once with PBS. Viability was assessed using the LIVE/DEAD Fixable Lime Viability Kit (L34991, Thermo Fisher; 100uL of 1:1000 dilution in PBS) for 20 minutes at 4°C, followed by two washes with PBS. Cells were then incubated with 2X CD16/CD32 Monoclonal Antibody Fc block (Thermo, Cat. #14-0161-86) at 1 μL/test in 50 uL Brilliant stain buffer (Thermo, Cat. #00-4409-75) for 20 minutes at 4°C. Without washing, 2X surface stain master mix was added directly to the Fc block for a final volume of 100 μL and incubated for 30 minutes at 4°C; CellBlox was included in the myeloid panel surface master mix. Cells were washed twice with staining buffer and centrifuged at 500xg for 5 minutes. For intracellular targets, cells were fixed following surface staining using IC Fix/Perm kit (Invitrogen, Cat. #88-8824-00) (Arginase-1) or FoxP3 Fix/Perm Buffer kit (Invitrogen Cat. #00-5523-00) (FOXP3) for 30 minutes at room temperature protected from light. Cells were centrifuged at 500xg for 5 minutes, washed once (twice for FOXP3) with 1X Permeabilization Buffer, and resuspended in approximately 100 μL Permeabilization Buffer. Directly conjugated intracellular antibodies were added and incubated for 30 minutes at room temperature, protected from light, followed by two washes in Permeabilization Buffer. Cell were fixed in 2% PFA in PBS overnight at 4°C. Before analysis, cells were pelleted at 500xg for 5 minutes and resuspended in flow cytometry staining buffer. Data acquisition was performed on an Attune NxT and analyzed using FlowJo V10.

### Histology

Following overnight fixation in 4% paraformaldehyde, samples were stored in 70% ethanol and then processed in an automated tissue processor (Leica TP1020), embedded in paraffin, and sectioned at 4-6 μm. Hematoxylin and Eosin (H&E) staining was performed according to standard procedures. Hematoxylin 560 (Leica Biosystems, Cat. #3801570) was filtered through Whatman #4 filter paper before use. Slides were deparaffinized in a 60°C oven for ∼1 hour, rehydrated through 3 washes of xylene (5 minutes each) and sequential ethanol washes (3 x 100%, then 95% and 70%, 2 minutes each), and rinsed in 3 changes of reverse osmosis (RO) water followed by 3 minutes in running RO water. Slides were incubated in hematoxylin for 4 minutes 30 seconds, rinsed in 3 changes of RO water, differentiated in Define MX (Leica Biosystems, Cat. #3803595) for 30 seconds, rinsed in running RO water for 2 minutes, dipped 20 times in Blue Buffer (Leica Biosystems, Cat. #3802915) over 30 seconds, and rinsed again in 3 changes of RO water with a final 3-minute running water wash. After a 2-minute incubation in 70% ethanol, slides were stained in Eosin 515 Phloxine (Leica Biosystems, Cat. #3801606) for 1 minute 30 seconds, dehydrated through 3 changes of 100% ethanol (2 minutes each), cleared in 3 changes of fresh xylene (2 minutes each), and mounted with CytoSeal XYL under #1.5 coverslips.

For mammary gland adipocyte analysis, H&E-stained sections were analyzed in fields of view within the mammary fat pad distal to the nipple and excluding epithelium. Individual adipocyte area was quantified by automated thresholding in NIH ImageJ.

Immunohistochemistry used the ABC kit (Vector Laboratories, Cat. #PK-6100) and DAB kit (Vector Laboratories, Burlingame, CA) per the manufacturer’s instructions. Antigen retrieval was done by boiling the slides in 10 mM sodium citrate for 15 minutes in a microwave. Primary antibodies included anti-cleaved caspase 3 (Cell Signaling, Cat. #9661S, 1:300), and anti-Ki67 (Cell Signaling, Cat. #12202, 1:400). Slides were counterstained with Hematoxylin QS (Vector Laboratories) and imaged at 20x using an ECHO Revolution slide scanner. Three fields of view per sample were analyzed, and the percentage of positive nuclei relative to total epithelial cells per duct was quantified manually. Four to six samples per genotype per age were assessed per cohort.

Immunofluorescence staining was used to assess mammary ductal organization in Cxcr2 WT, Het, and KO glands. Antigen retrieval was performed by boiling in 10 mM sodium citrate for 15 minutes in a microwave. Tissue sections were bordered with an ImmEdge hydrophobic barrier pen (Vector Laboratories, Cat. #H-4000) and blocked with 5% normal goat serum (Jackson ImmunoResearch, Cat. #005-000-121). Primary antibodies included anti-keratin-8 (Developmental Studies Hybridoma Bank, #TROMA-1-c, 1:500, 0.34 µg/mL), anti-keratin-14 (Covance, Cat. #PRB-155P-100, 1:5000, 0.2 µg/mL), and anti-smooth muscle actin (FITC-conjugated, Sigma, Cat. #F3777, 1:1000, 3 µg/mL). Secondary antibodies anti-rabbit Alexa Fluor 488 (Thermo Fisher, Cat. #A11008, 1:250, 8 µg/mL) and anti-rat Alexa Fluor 568 (Thermo Fisher, Cat. #A11077, 1:250, 8 µg/mL) were applied, and nuclei were counterstained with Hoechst 33342 (MilliporeSigma, Cat. #14533, 1:500, 1 μg/mL final) in blocking buffer for 1-1.5 hours at room temperature. After PBS washes, slides were coverslipped with Vectashield Vibrance Antifade mounting medium (Vector Laboratories, Cat. #H-1700) and imaged at 20x using an ECHO Revolution slide scanner with DAPI (ECHO, Cat. #CUBE6001), FITC (ECHO, Cat. #CUBE6002), and Texas Red (ECHO, Cat. #CUBE6004) filter sets.

Quantification of K8/K14 staining included manual counting of K8+, K14+, and K8+/K14+ cells as a percentage of total mammary epithelial cells per field of view, as well as mean fluorescence intensity and positive area per duct. Three duct images were chosen based on roughly same amount of epithelium per field of view and averaged per sample, and 2-4 samples per genotype were assessed. K8/SMA staining was evaluated qualitatively to confirm normal luminal and myoepithelial organization.

### Mammary gland whole mounts and carmine alum staining

Carmine alum staining solution was prepared by boiling 0.2% carmine (Sigma, Cat. #C1022) and 0.5% aluminum potassium sulfate (Sigma, Cat. #A7167) in distilled water for 20 minutes, filtering through a 0.22 μm membrane (Santa Cruz, Cat. #sc-258377), and adding a crystal of thymol (Sigma, Cat. #T0501) as preservative. Solution was stored at 4°C and reused until staining quality diminished.

The number 4 mammary glands from 3-, 5-, 8-, and 12-week-old virgin female Cxcr2 FVB/N mice were collected following CO_2_ euthanasia. Estrus stage was determined by vaginal cytology. Samples were compared based on estrus, with diestrus samples excluded and shown in supplemental data (none in 3-, 5-, and 8-week mice, three 12-week mice). Glands were spread on glass slides, fixed overnight in Carnoy’s solution (3:1 v/v ethanol:acetic acid), hydrated through a graded ethanol series, and stained overnight at 4°C in carmine alum solution. Tissues were dehydrated in an ethanol gradient, cleared in xylene for several days, and imaged at 1x and 2.5x on a Leica dissection microscope. Ductal area, length, and terminal end bud number were quantified in NIH ImageJ.

Ductal branching was analyzed using the Advanced Sholl Analysis plugin for Fiji (http://fiji.sc/Sholl_Analysis). Whole mount images were opened in ImageJ, and the mammary gland tissue outlined with the Freehand tool (“Edit > Clear Outside”). The lymph node was excised using the Cut tool. Color channels were split (“Image > Color > Split Channels”) and the blue channel, which provided optimal contrast, was selected for analysis. Background was subtracted (“Process > Subtract Background”) and noise removed (“Process > Noise > Despeckle”), with residual artifacts manually erased using the original image as reference. The image was binarized by thresholding (“Image > Adjust > Threshold”) to isolate mammary epithelium, and any gaps in epithelium were reconstructed using the Spray Can tool. A skeletonized image was generated (“Process > Binary > Skeletonize”), gaps filled by dilation (“Process > Binary > Dilate”), and the skeleton saved. Accuracy was verified by overlaying the skeleton on the original image at 30% opacity (“Image > Overlay > Add Image”).

For Sholl analysis, the skeleton was made binary (“Process > Binary > Make Binary”), and the scale was calibrated from the image scale bar (“Analyze > Set Scale”). The longitudinal growth radius was measured by drawing a line from the origin of the primary duct to the most distal epithelial point; the maximum across all images, rounded up to the nearest 0.5 mm, was used as the ending radius. Analysis was run via “Plugins > Advanced Sholl Analysis” with the following parameters: starting radius = 0.00 mm; radius step size = 0.1 mm; samples per radius = 1; integration = mean; enclosing radius cutoff = 1; primary branches = inferred from starting radius. Linear fit with best fitting degree was selected for unnormalized profiles, and the “Most Informative” normalization was selected for normalized profiles. The intersections mask was generated for each sample.

Primary endpoints were the number of radial intersections (N) and the Sholl regression coefficient (k), used to calculate branching density as ΣN / (MEA − LNA), where mean epithelial area (MEA) and lymph node area (LNA) were determined by manual outlining and “Analyze > Measure”. LNA was set to zero when epithelium had not reached the lymph node. Branching complexity (k) was derived from log(N/S) = −kr + m.

### Picrosirius red staining

Picrosirius red staining of paraffin-embedded mammary gland tissue sections from 5- and 8-week-old Cxcr2 WT, Het, and KO mice was used to quantify collagen content. Slides were deparaffinized in a 60°C oven for ∼1 hour followed by 2 washes in xylene (5 minutes each), rehydrated through 2 washes each of 100%, 95%, and 70% ethanol (3 minutes each) and 5 minutes in distilled water. Sections were counterstained in hematoxylin (Leica Biosystems, Cat. #3801570) for 30 seconds, rinsed in running tap water for 30 seconds, and incubated in Picrosirius staining solution (0.5 g Sirius Red [Fisher, Cat. #50-520-528] in 500 mL saturated picric acid [1.2% w/v; Ricca Chemical, Cat. #5860-16]) for 30-60 minutes. Slides were differentiated in two changes of acidified water (5 mL acetic acid per 1 L distilled water, 5 minutes each), dehydrated in 3 changes of 100% ethanol (3 minutes each), cleared in xylene, and mounted with CytoSeal XYL under #1.5 coverslips.

Slides were imaged in brightfield and polarized light on a Keyence BZ-X810 microscope. Polarized images were analyzed in ImageJ by converting to RGB stack and adjusting contrast and threshold to isolate collagen birefringence, using the brightfield image to restrict analysis to mammary gland tissue. Green birefringence was assigned to thin Type III collagen fibers and orange/red birefringence to thick Type I collagen fibers.

## Statistical analysis

Statistical analyses were performed in GraphPad Prism (Version 10.0). Overall survival was compared between groups using the log-rank test, with deaths censored for planned euthanasia or experimental endpoint sacrifice; pups that died prior to genotyping (∼3 weeks of age) were excluded. Body composition data from Cxcr2 WT, Het, and KO mice were compared by two-way ANOVA with multiple comparisons. Transplantation studies were analyzed using linear mixed-effects models in R, with stromal (recipient) and epithelial (donor) Cxcr2 genotype and their interaction as fixed effects and recipient mouse as a random effect to account for paired samples. Total branching intersection counts and adipocyte areas were log-transformed prior to analysis, while the remaining outcomes were analyzed on their original scales. Study was additionally included as a fixed effect in the adipocyte area model. Pairwise comparisons between WT and KO groups were assessed by Welch’s t-test or Mann-Whitney test as specified in figure legends. Results were considered significant at P<0.05.

## Acknowledgments

We would like to thank the members of the Littlepage lab for their helpful insights and comments throughout this study. We acknowledge the Notre Dame Integrated Imaging Facility (NDIIF) for imaging instrumentation and the Harper Cancer Research Institute and Tissue Core for instrumentation and support. We also thank the Freimann Life Sciences Center staff at the University of Notre Dame for the animal care and husbandry of mice used in this study.

## Statements and declarations

The authors have no competing interests to declare.

