## Supplemental Figure 1 for "Cxcr2 is required for ductal outgrowth and shapes the stromal microenvironment in postnatal mammary gland development"

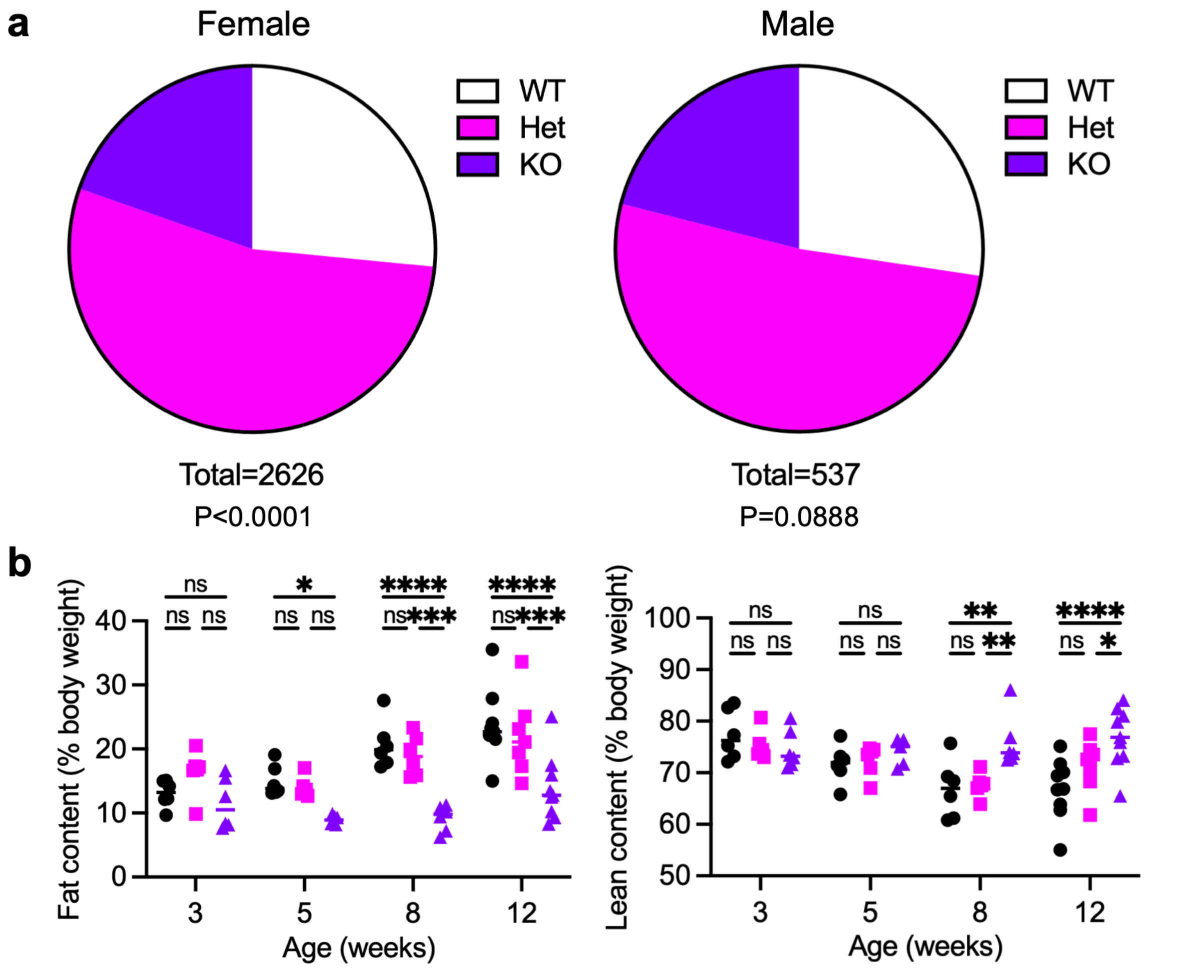


**Supplemental Fig. 1 Sex-specific genotype ratios and body composition in Cxcr2 KO mice.** (A) Genotype distribution in female (left; total = 2626) and male (right; total = 537) offspring from Cxcr2 Het × Het crosses compared to expected Mendelian ratios by chi-square test. (B) EchoMRI body composition analysis of fat content (left) and lean content (right), each as a percentage of body weight, in Cxcr2 WT, Het, and KO mice at 3, 5, 8, and 12 weeks (n = 6-9/genotype/age). Statistical analysis by two-way ANOVA with Tukey's multiple comparisons test. Lines indicate mean. ns, not significant; *p < 0.05, **p < 0.01, ***p < 0.001, ****p < 0.0001
