## Supplemental Figure 2 for "Cxcr2 is required for ductal outgrowth and shapes the stromal microenvironment in postnatal mammary gland development"

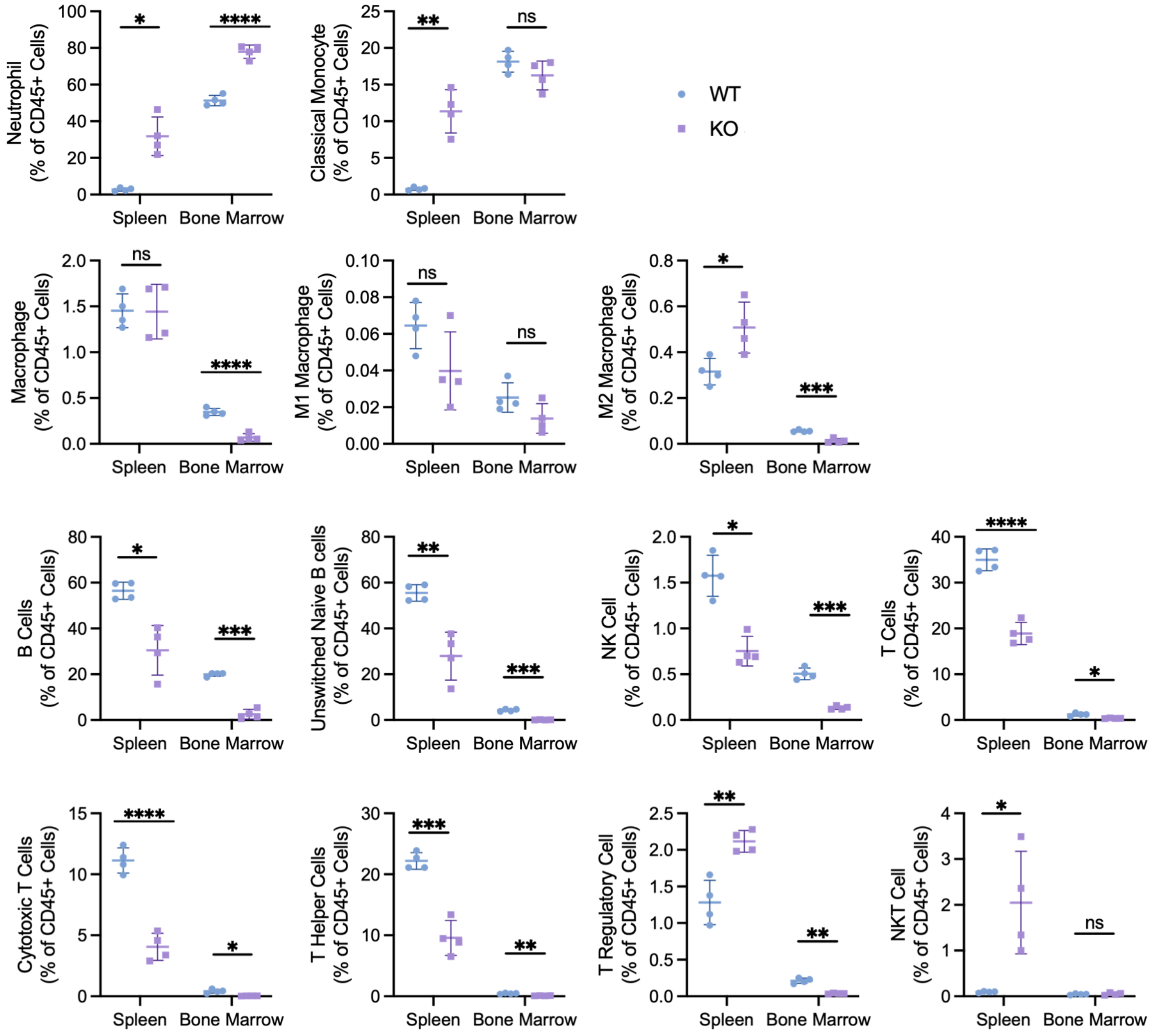


**Supplemental Fig. 2 Full flow cytometry profiling of spleen and bone marrow immune populations.** Spleen and bone marrow flow cytometry quantification of the full panel of immune populations in Cxcr2 WT and KO mice (n = 4/genotype): granulocytic and myeloid populations including neutrophils, PMN-MDSCs, monocytes, and M-MDSCs (as % of CD45+ cells); M1 and M2 macrophages (as % of CD45+ cells and as % of total macrophages); and lymphoid populations including B cells, unswitched naïve B cells, NK cells, total T cells, cytotoxic (CD8+) T cells, T helper (CD4+) cells, regulatory T cells, and NKT cells (as % of CD45+ or CD3+ cells as indicated on each axis). Population-defining gating markers are indicated in Table 2. Statistical analysis by Welch's t-test. Lines indicate mean. ns, not significant; *p < 0.05, **p < 0.01, ***p < 0.001, ****p < 0.0001
