## Supplemental Figure 3 for "Cxcr2 is required for ductal outgrowth and shapes the stromal microenvironment in postnatal mammary gland development"

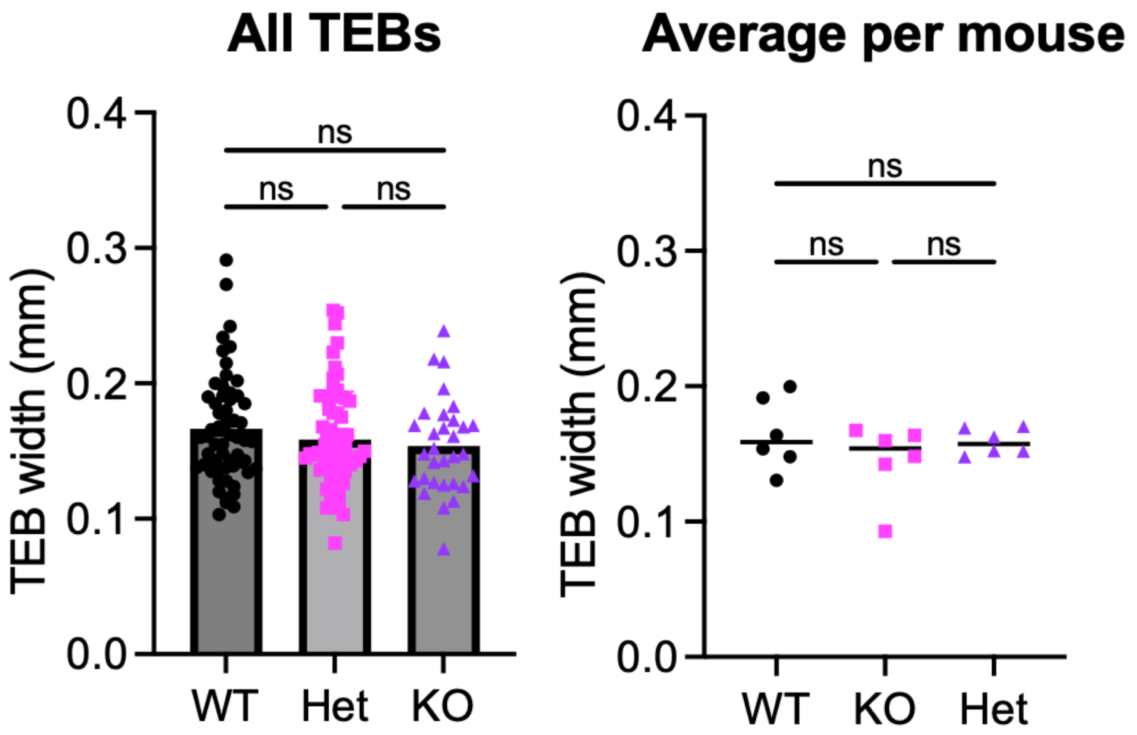


**Supplemental Fig. 3 Terminal end bud width does not differ by genotype.** (A) TEB width (mm) in Cxcr2 WT, Het, and KO 3-week mammary glands, shown as all measured TEBs pooled (left; n = 58 WT, 72 Het, 31 KO TEBs across 6 mice/genotype) and as the per-mouse average (right; n = 6 mice/genotype). Statistical analysis by one-way ANOVA with Tukey's multiple comparisons test. Lines indicate mean. ns, not significant
